# Design of a GAK/EGFR inhibitor set to interrogate the relationship of EGFR and GAK in chordoma

**DOI:** 10.1101/475251

**Authors:** Christopher R. M. Asquith, Kaleb M. Naegeli, Michael P. East, Tuomo Laitinen, Tammy M. Havener, Carrow I. Wells, Gary L. Johnson, David H. Drewry, William J. Zuercher, David C. Morris

## Abstract

We describe the design of a set of inhibitors to investigate the relationship between cyclin G associated kinase (GAK) and epidermal growth factor receptor (EGFR) in chordoma bone cancers. These compounds were characterized both *in vitro* and using in cell target engagement assays. The most potent chordoma inhibitors were further characterized in a kinome-wide screen demonstrating narrow spectrum profiles.

## Introduction

Chordomas are rare tumors arising along the bones of the central nervous system and spine with invasive and metastatic potential driven by the notochord transcription factor brachyury.^1^ Largely due to the sensitive tissues affected, treatment of chordoma is challenging. The first line of treatment is generally radical resection with complementary proton therapy. Recurrent or metastatic disease is often fatal, as surgical and chemotherapeutic options are limited. Improved chordoma therapies will require a deeper understanding of the oncogenesis of chordomas and the molecular biology of the tumors.^1,2^

One promising chemotherapeutic avenue in chordoma is inhibition of the epidermal growth factor receptor (EGFR).^3^ EGFR and its ligand, EGF, are highly expressed in chordomas, and copy number gains of EGFR occur in 40% of chordomas.^3^ Compounds that inhibit EGFR have shown efficacy against independently-derived chordoma cell lines and murine xenograft models.^4,5^ These results have led to clinical evaluation of the irreversible EGFR inhibitor afatinib for patients with EGFR-expressing tumors.^4^ EGFR inhibitors are not expected to be effective against all chordomas. For example, one patient-derived chordoma cell line overexpresses EGFR but exhibits marked resistance to EGFR inhibitors due to high levels of activated MET.^6^ Hence there is need for chemotherapeutic targets beyond EGFR.

EGFR inhibitors based on the anilino-quin(az)oline scaffold often carry concurrent cyclin G associated kinase (GAK) inhibitory activity, and modulation of either of these kinases may affect the expression/activity of the other.^7,8^ The quinazoline-based clinical kinase inhibitors gefitinib, erlotinib, afatinib, and vandetanib have activity in chordoma cell lines (**Figure 1**).^9^ These drugs were designed to target EGFR and show similar or higher affinity for GAK and several other kinases, making them ineffective tools for direct integration of EGFR and GAK biology. We were interested in separation of the activities of both GAK and EGFR to assess the potential of GAK as a therapeutic target for chordoma either as a single target or in combination with EGFR inhibition.

**Figure 1.**
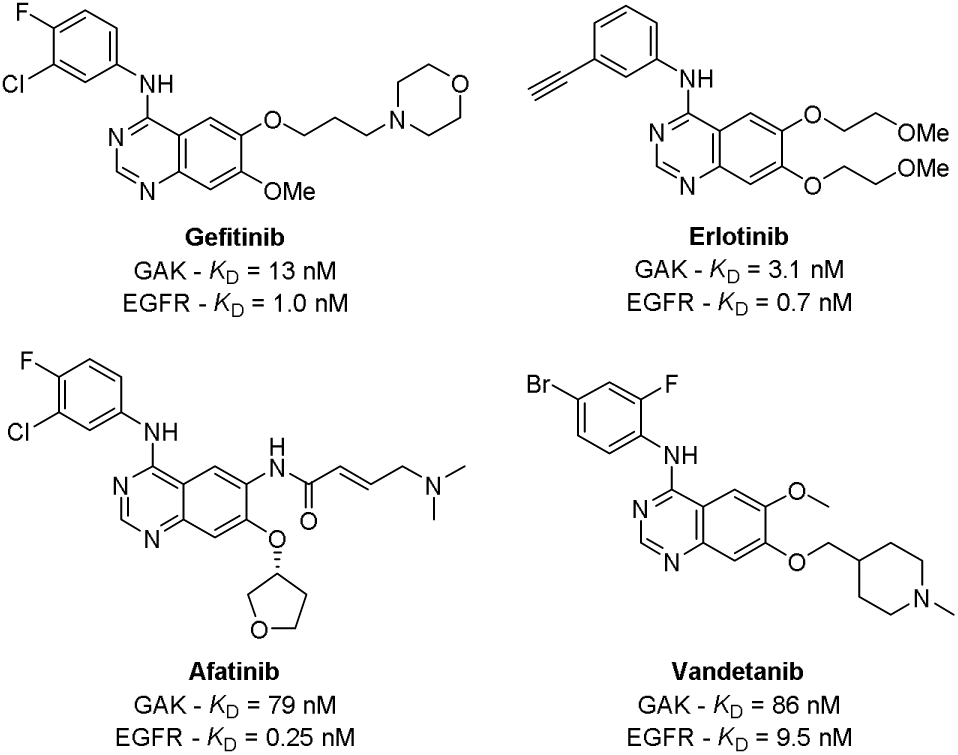
Previously reported inhibitors of GAK and EGFR.

## Results

We first selected a set of literature inhibitors that covered a cross section of quinoline/quinazoline EGFR chemical space (**Figure S1**). The compounds were screened directly against two chordoma cell lines, U-CH1 and U-CH2 (**Table 1**), the latter of which is notably resistant to EGFR inhibitors.^6^ This enabled us to identify several kinases including GAK as targets of interest, with EGFR and GAK the most prevalent across the series of inhibitors (**Table S1**). EGFR inhibitors utilizing the 4-anilino-quin(az)olines are often associated with off-target GAK activity.^9-15^ There was not a linear relationship between EGFR and/or GAK inhibition and anti-proliferative effects.

**Table 1.**
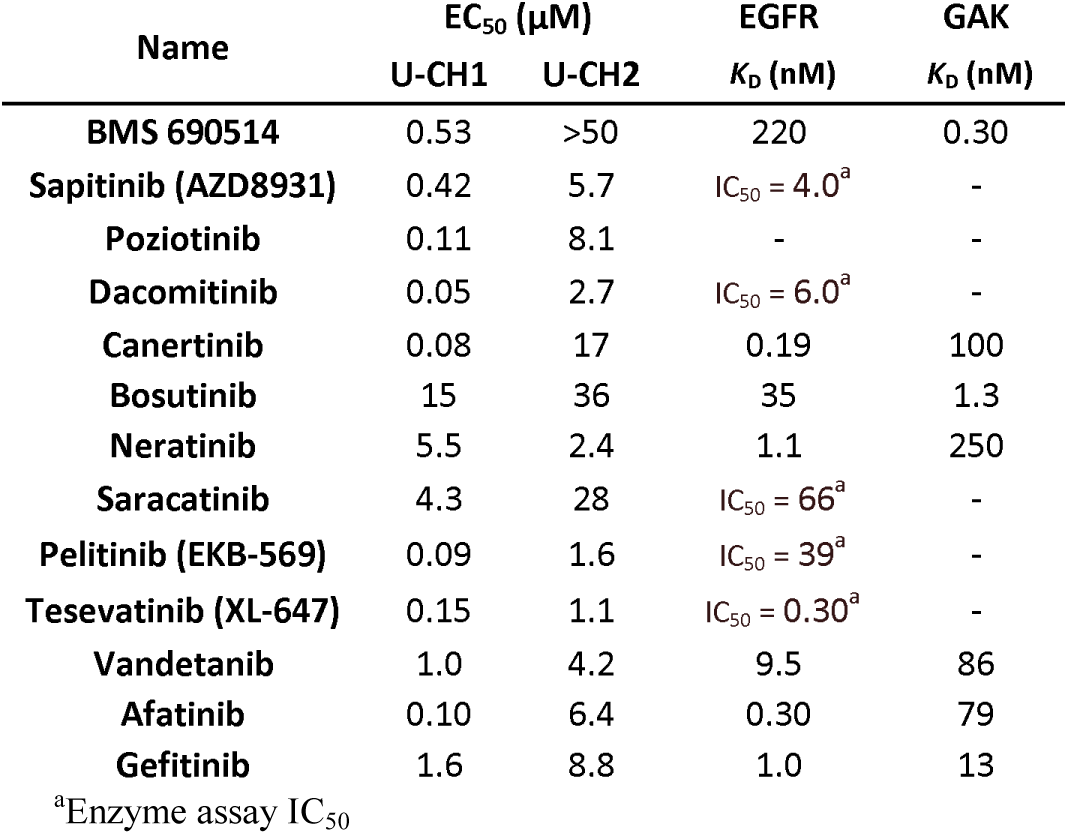
Literature inhibitors screening and reported activities

The different and sometimes broad kinome inhibition profiles possessed by each inhibitor was a confounding factor in establishing the molecular basis for the cellular activity. There was a clear trend between inhibiting GAK or EGFR and effects on the U-CH1 cell line. Generally, potent EGFR inhibition led to a strong response in U-CH1 with a generally weaker response in U-CH2. The effects of GAK inhibition were less clear, but BMS-690514 showed moderate potency against U-CH1 with relatively weak EGFR activity and a potent GAK profile with a >700-fold window over EGFR, while this does not rule out an EGFR contribution this would limit the effect.^10^ BMS-690514 had 5 other kinases with *K*_D_ < 225 nM, including RET, ABL2, SIK2, ABL1, PRKD2, and TNK2. The inhibition of any of these kinases, alone or in combination, could be responsible for the growth phenotype observed in U-CH1, confounding the interpretation of the result.^16^

We sought to determine the effect of both GAK and EGFR inhibition utilizing the 4-anilino-quin(az)oline scaffold. Molecular modelling of the scaffold in GAK and EGFR (**Figure 2A-D**) showed an ability to increase selectivity for EGFR over GAK by utilizing the deeper hydrophobic pocket of EGFR not present in GAK.

**Figure 2.**
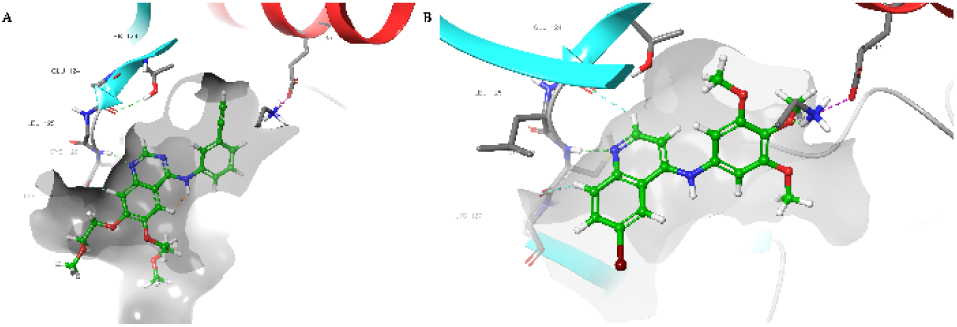
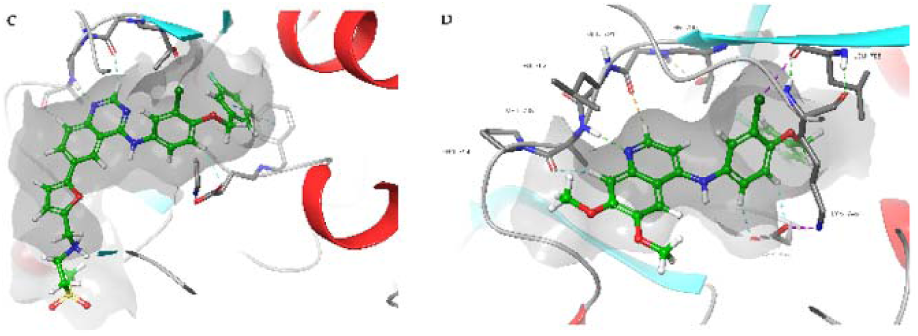
Design strategy for GAK/EGFR inhibitors: (A) Erlotinib and (B) 3 docked in GAK and (C) lapatinib and (D) **15** docked in EGFR demonstrating the hydrophobic pocket. (see SI for detailed images)

We synthesized compounds **1**-**19** through nucleophilic aromatic displacement of 4-chloroquin(az)olines (**Scheme 1**) to furnish the products in good yields (52-89%). The only low yielding analog was the trifluoro quinazoline (**2**) which was consistent with previous reports and likely arose from attenuation of aniline nucleophilicity by the powerful electron withdrawing effect of the trifluoro group.^17^

**Scheme 1.**
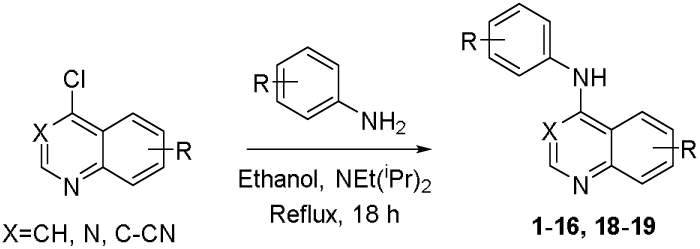
General synthetic procedure

We designed the compounds to exploit the structure activity relationships between GAK and EGFR within the quin(az)oline scaffold (**Table 2**). Compound **1** was previously identified as a narrow spectrum inhibitor of GAK with only three other off-targets with *K*_D_ < 1 µM in a kinome wide screen and a negligible affinity for EGFR at 1 μM.^17-19^ The corresponding quinazoline (**2**) had a 6-fold drop off *in vitro* and a 26-fold drop off in cells against GAK but a corresponding increase in EGFR biochemical activity.^17^ Compound **3**, in which the triflurormethyl of **1** was replaced by bromine, was highly selective, with only RIPK2 as an off-target across the kinome with a small increase in GAK cellular potency.^18^ Previous reports suggested the activity of the quin(az)oline can be independently modulated for both GAK and EGFR while maintaining a narrow spectrum profile across the kinome. The switch to the 6,7-dimethoxy substitution (**4**) showed potent GAK activity both biochemically and in cells.^17^ However **4** possessed previously uncharacterized EGFR activity albeit with a 140-fold window *in vitro* and only limited cell activity. The quinazoline (**5**) and 3-cyanoquinoline (**6**) both showed improved potency with both GAK and EGFR with **6** showing low single digit nanomolar potency in cells against GAK. **6** is the most potent GAK inhibitor reported to date.

**Table 2.**
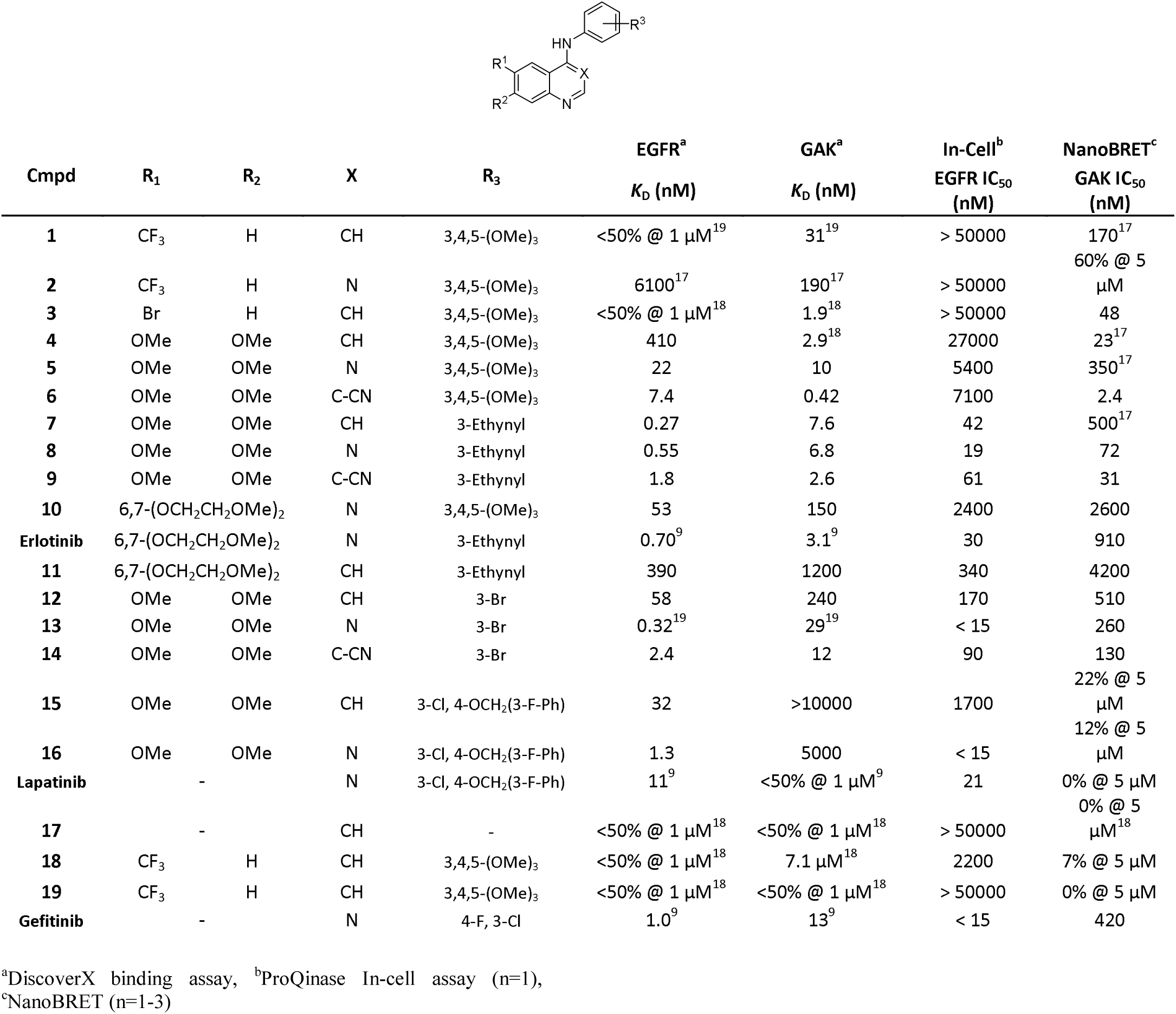
GAK and EGFR *in vitro* and in cell biochemical characterization.

The switch to the *meta*-acetylene (**7**-**9**) across the same range of hinge binder maintained GAK activity but significantly increased the potency of cellular target engagement against EGFR by more than 100-fold across the three compounds. This trend was mirrored when extending the 6,7-dimethoxy substitution to the 6,7-dimethoxyethoxy seen in erlotinib. The matched pairs of **10**/erlotinib and **11**/erlotinib demonstrate a well of activity around the quinazoline of erlotinib. We observed a significant decrease in EGFR inhibition with the switch from quinazoline to quinoline which is consistent with most matched pairs in the series and consistent with a previous report on EGFR inhibition for **11**/erlotinib.^20^ However, the drop in GAK activity between erlotinib and **11** was more surprising and suggested a more complex structure activity relationship when extending beyond small substitutions on the quino(az)oline ring system. Surprisingly erlotinib had a very activity both biochemically and in cells. The EGFR activity in this series increased with cellular potency approaching single digit nanomolar for **13**. This result demonstrated that GAK activity of the anilino-quino(az)oline scaffold can be reduced while maintaining potency on other kinases. We then sought to exploit the lapatinib scaffold to remove the GAK activity (Figure 2). We designed **15** and **16** using the lapatinib aniline portion as the ‘head group’ while having a simple 6,7-dimethoxy substituted quinoline (**15**) or quinazoline (**16**). These compounds demonstrated very limited GAK activity but moderate potency on EGFR both biochemically and in cells. We also included several control compounds (**17**-**19**) to counteract the off-target RIPK2 activity associated with this series (**17**) and the controls (**18**-**19**) that showed only limited activity.^18^ Gefitinib showed an activity profile similar to erlotinib and lapatinib in both UCH-1 and UCH-2. We then screened weak GAK cellular engagement despite a single digit nanomolar *K*_D_.

A switch to a *meta*-bromo (**12**-**14**) to increase the occupancy of the *meta*-acetylene led to an observed decrease in GAK these compounds using two clinically relevant chordoma cell lines, with different clinical presentations (**Table 3**).

**Table 3.** Chordoma and Fibroblast screening results

| Cmpd | U-CH1 | U-CH2 | WS1 |
| --- | --- | --- | --- |
| | $EC_{50} (\mu M)^a$ | | |
| **1** | >50 | 33 | 21 |
| **2** | 50 | >50 | 45 |
| **3** | >50 | >50 | 11 |
| **4** | 11 | 25 | 14 |
| **5** | 7.3 | >10 | 28 |
| **6** | 11 | >10 | >50 |
| **7** | 2.1 | >50 | 7.7 |
| **8** | 0.92 | >50 | 4.4 |
| **9** | 17 | >50 | >50 |
| **10** | 25 | >50 | >50 |
| Erlotinib | 5.1 | 18 | 8.6 |
| **11** | 7.6 | 16 | 12 |
| **12** | 3.6 | 7.9 | 2.9 |
| **13** | 0.66 | 26 | 9.7 |
| **14** | 8.8 | >50 | 14 |
| **15** | 2.6 | 2.9 | 8.6 |
| **16** | 13 | >50 | >10 |
| Lapatinib | 3.2 | >50 | 13 |
| **17** | >50 | 30 | >50 |
| **18** | >50 | >50 | >50 |
| **19** | 37 | >50 | 49 |
| Gefitinib | 1.4 | 23 | 23 |

<sup>a</sup>Standard mean in (n=1-3 see SI for errors)

We first screened the compounds containing the trimethoxy aniline (**1**-**3**) with in-cell GAK potency and no in-cell EGFR activity and found only a hint of inhibition on U-CH2 cell line with **1**. Increasing the potency on GAK with a change in substitution to 6,7-dimethoxy and a range of hinge binders (**4**-**6**) yielded a boost in potency on U-CH1 by more than 5-fold to single digit micromolar for **5**, without increased toxicity in normal human skin fibroblast (WS1) cells.

The *meta*-acetylenes (**6**-**9**) yielded a further potency boost, potentially due to the corresponding increase in EGFR activity. **7** and **8** showed some WS1 inhibition, with no activity on U-CH2. Interestingly the 3-cyano quinoline matched pair derivative (**9**) showed no toxicity on WS1 consistent with compound **6**. Combining the erlotinib quinazoline fragment with the trimethoxy aniline (**10**) showed limited activity on all cell lines. Erlotinib and the erlotinib quinoline derivative (**11**) had activity on all cell lines with some WS1 toxicity. However, there was only limited improvement in potency relative to **4** for both U-CH1 and U-CH2 and no improvement compared to both **7** and **8** for U-CH1.

Installation of the *meta*-bromine on the 6,7-dimethoxy core (**12**-**14**) increased EGFR activity and proved to be the most effective set of compounds tested with the matched pair **12** and **13** showing that the quinoline (**12**) is favored for U-CH2 growth inhibition, and the quinazoline (**13**) is favored for U-CH1. **14** was considerably less active than **12** and **13**. Compound **13** was most potent compound tested against U-CH1 (EC_50_ = 0.66 μM), but the more striking result was the EGFR inhibitor-resistant cell line U-CH2 showed relatively high sensitivity to **12**.

The two lapatinib derivatives **15** and **16** containing the lapatinib aniline and the 6,7-dimethoxy core were designed to remove the influence of GAK inhibition from any phenotypic response. The quinoline (**15**) showed the highest potency against U-CH2 (EC_50_ = 2.9 μM) with only limited toxicity. However, this compound showed weak EGFR activity in cells, confounding target identification. In contrast, **16** showed high cellular EGFR potency and only activity against U-CH1. As previously reported, lapatinib had low micromolar activity against U-CH1 and no activity against U-CH2^6^ We also observed a small amount of associated WS1 toxicity. The RIPK2 control compound **17** with no GAK or EGFR activity had limited inhibition as did **18** and **19**.^17^ Gefitinib was the only literature compound to show some activity on U-CH2 with a comparable profile to lapatinib on the other cell lines.

The compounds that showed the highest efficacy against the U-CH2 cell line were then evaluated against five further patient-derived chordoma cell lines (**Table 4**). The 5 cells lines showed varying inhibitor sensitivity, but all appeared less responsive

Compound **4** showed moderate activity across all cell lines consistent with the matched pair quinoline **7**. The switch to a quinazoline (**8**) produced quite a different profile with no activity against CH22, U-CH12 and U-CH14. Erlotinib was inactive up to 50 μM against all cell lines. The matched pair quinoline (**12**) displayed a similar profile to **8** with no activity against U-CH12 and U-CH14 but also U-CH7; with and sensitivity against UMChor1 and CH22, with only UMChor1 consistent between both **8** and **12**. Compound **15** showed single digit micromolar activity across all five cell lines and U-CH1 and U-CH2, with lapatinib showing a general >10-fold drop off in potency across the panel relative to U-CH1. Gefitinib only showed activity on the apparent most sensitive cell lineUMChor1, despite moderate activity on U-CH1 and U-CH2.

Having established activity in U-CH1 and U-CH2 cell lines, we next wanted to investigate downstream effects on EGFR (**Figure 3**) Short-term compound treatment with **7** or **8** in U-CH1 and U-CH2 cells did not affect total EGFR levels; however, phosphorylated EGFR was reduced, indicative of reduced EGFR signaling, a phenotype consistent with the action of other EGFR inhibitors in chordoma.^4-6^

**Figure 3.**
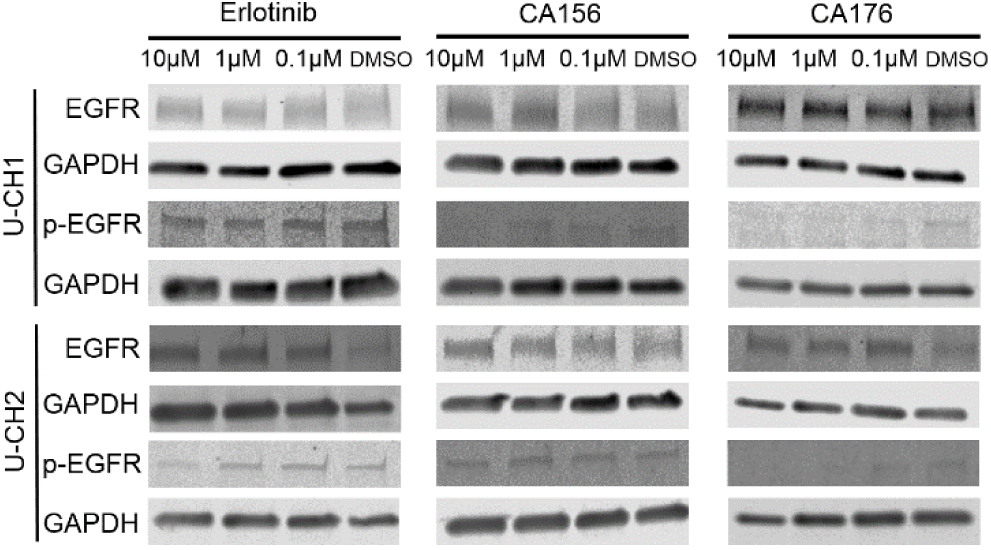
Western blot of EGFR and p-EGFR (Y1068) following erlotinib, **7**(CA156) and **8** (CA176) treatment

**Table 4.** Investigation of across 5 chordoma cell lines

| Name | U-CH7 | UMChor1 | CH22 | U-CH12 | U-CH14 |
| --- | --- | --- | --- | --- | --- |
| | $EC_{50} (\mu M)^a$ | | | | |
| **4** | 30 | 24 | 12 | 33 | 19 |
| **7** | 20 | 14 | 13 | 30 | 13 |
| **8** | 23 | 18 | >50 | >50 | >50 |
| Erlotinib | >50 | >50 | >10 | >50 | >50 |
| **12** | >50 | 17 | 6.5 | >50 | >50 |
| **15** | 2.7 | 1.5 | 1.0 | 5.5 | 7.0 |
| Lapatinib | 26 | 25 | 25 | 33 | >50 |
| Gefitinib | >50 | 20 | >50 | >50 | >50 |

<sup>a</sup>Standard mean (n=1-3 see SI for errors)

Compound **4** showed a narrow spectrum kinome profile, only GAK and RIPK2 were bound as assessed by multiplexed kinase inhibitor bead set (MIBS) and quantitative mass spectrometry (MS) detection of kinase peptides.^17^ The kinome selectivity profile of the other lead compounds (**7**, **8**, **12 & 15**) displayed narrow spectrum kinome profiles on >300 wild type kinases (**Supp. Info. 14.1.**). Compound **7** bound GAK, RIPK2, NEK1 and STRDA, while **8** associated only with GAK and limited EGFR. Compound **12** had binding to GAK, EGFR, PKM and BUB1, while **15** showed binding to EGFR, AURKB, MAP2K5 and NME2.

## Discussion

Chordoma is a difficult cancer to treat due to the slow growing nature and the integrated microenvironment surrounded by delicate tissues around the tumor. Chordomas have a high level of variance in presentation and susceptibility to treatment. This heterogeneity precludes a general approach to treatment and necessitates the need for personalization. EGFR inhibitors have been used to target chordomas, but the variation in efficacy across inhibitors is not presently understood and likely driven by off-target effects working additively or synergistically with EGFR inhibition. Conversely the off-tareget kinases could also be limiting efficacy.

GAK is unlikely to be one of these targets with compound **3** and the corresponding controls (**18 & 19**) showing limited effect. However, as EGFR inhibition is increased the efficiency of the compounds also increases. Compound **13** is the most potent compound ever tested against U-CH1 which is likely due to high in-cell potency on EGFR. The matched pair compound **12** was a more significant result with increased potency on the U-CH2 cell line relative to **13**. However, compound **15**, a derivative of lapatinib demonstrated the most diverse inhibition profile across patient derived cell lines with good selectivity over normal human skin fibroblast (WS1) toxicity. Interestingly we observed that a switch between the quinazoline (**8**) and quinoline (**7**) produced a radically different profile with no activity against CH22, U-CH12, and U-CH14, highlighting the complexity of chordoma biology.

This set of compounds provides a useful tool for the interrogation of GAK and EGFR and enables multi-point separation of their phenotypes. Profiling of the compounds with MIBS revealed that the compounds have a narrow kinome profile with GAK and EGFR likely the principal components of activity and biology observed.

## Experimental

### Biology & Screening

Chordoma cell lines were cultured as described previously (See SI for details). Screening was performed on multiple platforms (See SI for details). ^6,17^

### MIBS

Kinome profiling was performed as previously described (See SI for details).^17^

### Computational Modelling

was performed using Schrödinger Maestro software (See SI for details).

### Chemistry

**General procedure for the synthesis of 4-anilinoquinolines**: 6-Bromo-4-chloroquinoline (1.0 eq.), 3,4,5 trimethoxyaniline (1.1 eq.), and ^i^Pr_2_NEt (2.5 eq.) were suspended in ethanol (10 mL) and refluxed for 18 h. The crude mixture was purified by flash chromatography using EtOAc:hexane followed by 1-5 % methanol in EtOAc. After solvent removal under reduced pressure, the product was obtained as a free following solid or recrystallized from ethanol/water. Compounds (**1**-**2**, **4**-**5**, **7**, **13**)^17^; (**3**, **18**-**19**)^18^ **17** was purchased from MedChemExpress, (Monmouth Junction, NJ, USA). Compounds **6**, **9**-**11**, **13**-**14** and **16** data can be found in the SI.

### *N*-(3-ethynylphenyl)-6,7-dimethoxyquinazolin-4-amine (**8**)

colorless solid (181 mg, 0.594 mmol, 89 %) MP 237-239 ^o^C; ^1^H NMR (500 MHz, DMSO-d_6_) δ 11.48 (s, 1H), 8.85 (s, 1H), 8.38 (s, 1H), 7.88 (t, J = 1.8 Hz, 1H), 7.79 (ddd, J = 8.1, 2.2, 1.1 Hz, 1H), 7.49 (t, J = 7.8 Hz, 1H), 7.40 (dt, J = 7.7, 1.3 Hz, 1H), 7.37 (s, 1H), 4.28 (s, 1H), 4.02 (s, 3H), 3.99 (s, 3H). ^13^C NMR (125 MHz, DMSO-d_6_) δ 158.1, 156.3, 150.2, 148.8, 137.4, 136.0, 129.2, 129.2, 127.6, 125.3, 122.0, 107.4, 104.1, 99.9, 82.9, 81.3, 57.0, 56.5. HRMS m/z [M+H]^+^ calcd for C_18_H_16_N_3_O_2_: 290.1293, found 306.1230, LC t_R_ = 3.41 min, >98% Purity.

### N-[3-chloro-4-(3-fluorophenoxymethyl)phenyl]-6,7-dimethoxyquinolin-4-amine (**15**)

colorless solid (200 mg, 0.456 mmol, 68 %) MP 244-246 ^o^C; ^1^H NMR (400 MHz, DMSO-d_6_) δ 10.78 (s, 1H), 8.32 (d, J = 6.9 Hz, 1H), 8.19 (s, 1H), 7.62 (d, J = 2.5 Hz, 1H), 7.61 – 7.21 (m, 6H), 7.26 – 7.11 (m, 1H), 6.64 (d, J = 6.9 Hz, 1H), 5.31 (s, 2H), 4.00 (s, 3H), 3.96 (s, 3H). ^13^C NMR (101 MHz, DMSO-d_6_) δ 162.2 (d, J = 243.7 Hz), 154.5, 153.3, 152.2, 149.4, 139.8, 139.3 (d, J = 7.6 Hz), 135.2, 131.1, 130.6 (d, J = 8.4 Hz), 127.4, 125.7, 123.4 (d, J = 2.7 Hz), 122.2, 115.0, 114.8 (d, J = 20.9 Hz), 114.1 (d, J = 21.9 Hz), 111.5, 102.8, 99.8, 99.1, 69.5 (d, J = 1.9 Hz), 56.8, 56.1. HRMS m/z [M+H]^+^ calcd for C_24_H_21_N_2_O_3_ClF: 439.1225 found = 439.1213; LC t_R_ = 5.28 min, >98% Purity.

0000-0001-5871-3458 (Christopher Asquith - Orcid)

0000-0002-7551-4108 (Kaleb Naegeli – Orcid)

0000-0001-5973-5798 (David Drewry – Orcid)

0000-0002-9836-0068 (William Zuercher - Orcid)

0000-0002-2910-7544 (David Morris – Orcid)

## ASSOCIATED CONTENT

### Supporting Information

This material is available free of charge via the Internet at http://pubs.acs.org.

## AUTHOR INFORMATION

## Author Contributions

The manuscript was written through contributions of all authors. All authors approved of the final version of the manuscript.

## Notes

The authors declare no competing financial interests.

## ACKNOWLEDGMENT

The SGC is a registered charity (number 1097737) that receives funds from AbbVie, Bayer Pharma AG, Boehringer Ingelheim, Canada Foundation for Innovation, Eshelman Institute for Innovation, Genome Canada, Innovative Medicines Initiative (EU/EFPIA) [ULTRA-DD grant no. 115766], Janssen, Merck KGaA Darmstadt Germany, MSD, Novartis Pharma AG, Ontario Ministry of Economic Development and Innovation, Pfizer, São Paulo Research Foundation-FAPESP, Takeda, and Wellcome [106169/ZZ14/Z]. The UNC Catalyst for Rare Diseases is supported by UNC-CH and Eshelman Institute for Innovation. We thank Biocenter Finland/DDCB for financial support and the CSC-IT Center for Science Ltd. (Finland) for allocation of computational resources and Prof. Antti Poso (University of Eastern Finland) is thanked for informative discussions. We are grateful to the Chordoma Foundation for providing the cell lines and Dr. Brandie Ehrmann for LC-MS/HRMS support provided by the Mass Spectrometry Core Laboratory at the University of North Carolina at Chapel Hill. We also thank the team including Dr. Alokta Chakrabarti and Dr Katrin Schlie at ProQinase for their assistance and advice.

---

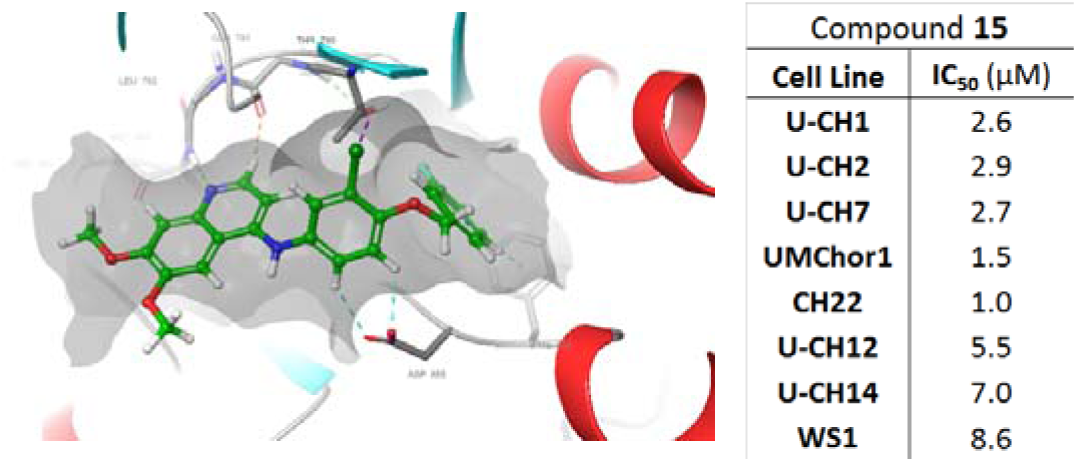

